# Click-train evoked steady state harmonic response as a novel pharmacodynamic biomarker of cortical oscillatory synchrony

**DOI:** 10.1101/2023.07.17.549322

**Authors:** Deepshila Gautam, Muhammad Ummear Raza, M Miyakoshi, JL Molina, YB Joshi, PE Clayson, GA Light, NR Swerdlow, Digavalli V. Sivarao

## Abstract

Sensory networks naturally entrain to rhythmic stimuli like a click train delivered at a particular frequency. Such synchronization is integral to information processing, can be measured by electroencephalography (EEG) and is an accessible index of neural network function. Click trains evoke neural entrainment not only at the driving frequency (F), referred to as the auditory steady state response (ASSR), but also at its higher multiples called the steady state harmonic response (SSHR). Since harmonics play an important and non-redundant role in acoustic information processing, we hypothesized that SSHR may differ from ASSR in presentation and pharmacological sensitivity. In female SD rats, a 2 s-long train stimulus was used to evoke ASSR at 20 Hz and its SSHR at 40, 60 and 80 Hz. Narrow band evoked responses were evident at all frequencies; signal power was strongest at 20 Hz while phase synchrony was strongest at 80 Hz. SSHR at 40 Hz took the longest time (∼180 ms from stimulus onset) to establish synchrony. The NMDA antagonist MK801 (0.025-0.1 mg/kg) did not consistently affect 20 Hz ASSR phase synchrony but robustly and dose-dependently attenuated synchrony of all SSHR. Evoked power was attenuated by MK801 at 20 Hz ASSR and 40 Hz SSHR only. Thus, presentation as well as pharmacological sensitivity distinguished SSHR from ASSR, making them non-redundant markers of cortical network function. SSHR is a novel and promising translational biomarker of cortical oscillatory dynamics that may have important applications in CNS drug development and personalized medicine.

## 1. Introduction

Cortical oscillations, recorded through EEG, impact how the brain processes information (Buzsaki and Draguhn, 2004). For example, the effect of a discrete sensory stimulus on a cortical network’s response can vary substantially in relationship to the on-going phase of the cortical field oscillation (Haegens and Zion Golumbic, 2018; Lakatos et al., 2019). Experimentally, neural networks can be entrained by rhythmic sensory stimuli like light flashes or click trains delivered at a particular driving frequency (F) (Coffey et al., 2019; Gorina-Careta et al., 2019; Norcia et al., 2015). In schizophrenia, endogenous and evoked neural synchrony is disrupted, especially at the gamma frequency of ∼ 40 Hz (Brenner et al., 2009; Kwon et al., 1999; Uhlhaas and Singer, 2010). This can be readily demonstrated via EEG entrainment to 40 Hz click trains as measured by the auditory steady state response (ASSR) (Brenner et al., 2009; Kwon et al., 1999; Light et al., 2006). Although less studied, disruptions at driving frequencies other than 40 Hz have also been noted (Hamm et al., 2011; Krishnan et al., 2009; Puvvada et al., 2018).

An advantage of ASSR as a measure of neural synchrony vs. other types of task-driven gamma oscillations is that the former lends itself to excellent duration control and rapid sampling and demands little conscious effort or intent from subjects. Furthermore, while the evoked signal is centered on a narrow frequency (F) and its multiples (see below), the ongoing neural activity reflected as the “background” EEG signal tends to be in the broad band, giving ASSR an outstanding signal/noise advantage (Norcia et al., 2015). Since this stimulus-driven neural entrainment is well conserved in mammals, ASSR is an attractive translational tool for demonstrating functional engagement and for dose selection in CNS drug development studies.

Complex sounds like clicks at a particular rate carry signal energy not only at that F but also at its integer multiples (n^*^F), called “harmonics.” In acoustic processing of sound, owing to their frequency difference, F and n^*^F trains activate different regions of the cochlear basilar membrane and are routed and processed somewhat differently (Oxenham, 2018). In terms of psychophysics, harmonics contribute to the distinctness of complex sounds (Wang, 2013). An important reason why the same musical note with the same fundamental frequency played by two different musical instruments sounds distinct is because of the difference in their harmonics. Although only multiples of F, harmonics play a non-redundant role in sound processing (Oxenham, 2012; Plack et al., 2006). For example, when the fundamental frequency of a musical note is somehow occluded, the brain can still perceive it by interpolating from the harmonics, an ability that is evolutionarily conserved across species (Hoeschele, 2017; Schouten and Ritsma, 1962). Harmonics in speech improve its intelligibility in situations of high ambient noise (McPherson et al., 2021).

While some data exist to indicate that steady state stimuli-driven harmonic responses may be disrupted in mental illness (Jin et al., 2000; Rice et al., 1989; Spencer et al., 2009), there has not been a focus on this aspect of the steady state response until recently (Molina et al., 2022; Raza and Sivarao, 2021a). One reason could be an assumption that harmonics may not carry information that is novel, beyond what is already captured by the ASSR. Indeed, multiple reports have demonstrated the existence of click train-driven harmonics in their data but did not explicitly address or discuss these (Ogyu et al., 2023; Tsuchimoto et al., 2011; Zhou et al., 2018). While few studies examine ASSR and harmonics, there have been reports that showed ASSR at 20 Hz (F) was augmented or unaffected in schizophrenia patients while its 2^nd^ harmonic at 40 Hz was disrupted (Kwon et al., 1999; Ogyu et al., 2023; Spencer et al., 2008; Vierling-Claassen et al., 2008). It is unknown whether higher harmonics of 20 Hz were present in these studies (e.g., 60 or 80 Hz) and if so, whether they were also disrupted.

Neurophysiological deficits similar to those exhibited by schizophrenia patients are experimentally reproduced in healthy human subjects, as well as in preclinical species, by acute administration of NMDA receptor antagonists (Curic et al., 2019; Lazarewicz et al., 2010; Lee and Zhou, 2019; Raza and Sivarao; Sivarao et al., 2016; Sivarao et al., 2013). In this report, we used a 20 Hz click train as a probe stimulus in female SD rats and examined the ASSR at 20 Hz and at the first three narrow band SSHR (∼ 40, 60 and 80 Hz). We then compared the sensitivity of the ASSR and SSHR to graded doses of a high affinity NMDA receptor antagonist, MK801. Since SSHR may result from activating a different neural network from the one generating the response at F (Hamm et al., 2012; Heinrichs-Graham and Wilson, 2012; Pastor et al., 2007), we hypothesized that that there would be measurable differences in the characteristics of SSHR and ASSR and in their responses to drug challenge.

## 2. Methods

### 2.1. General

All experimental procedures were approved by the Institutional Animal Care and Use Committee of the East Tennessee State University. Six-eight week-old female Sprague Dawley rats (N = 12; 150–200 g; Envigo (Indianapolis, IN)) were group housed with food and water <u>ad lib</u>.

### 2.2. Surgical

All surgeries were performed as previously described (Raza and Sivarao, 2021b). In brief, animals were anaesthetized with isoflurane (induction 5%; maintenance, 1-2% in 1 L/min oxygen) and secured in a stereotaxic frame. Alcohol and povidone-iodine swabs were used to clean the area. Bupivacaine (0.25 %, 8mg/kg, sc) was infiltrated in the subcutaneous field for local anesthesia and ketoprofen (5 mg/kg, sc) was used for perioperative analgesia. A midline incision was made, and soft tissue was removed through blunt dissection to expose the skull. A solution of 3% hydrogen peroxide was introduced drop by drop to disinfect the area and was mopped up and allowed to dry. A frontal electrode (2 mm anterior and 1 mm lateral to bregma) was used for recording the ASSR while two electrodes placed above the occipital bone served as ground and reference (2 mm posterior to lambda and 2 mm on either side from midline). The frontal electrode targeted the underlying secondary motor cortex, considered a part of the rodent prefrontal cortical network, based on thalamic input and function (Barthas and Kwan, 2017). Two other electrodes were placed, one on the vertex and another on the primary auditory cortex. However, these data have not been analyzed at this time. All electrodes with attached wires were carefully screwed into the burr holes while keeping the underlying dura intact. Electrodes were poured over with dental cement slush. A recording head mount (Pinnacle Technology, Lawrence, KS) was placed on the dental cement after it had cured, and electrode wires were soldered to the head mount leads. Exposed hardware was covered with a second layer of dental cement. The surgery area was closed with sutures and rats were given at least 10 days to recover with daily enrofloxacin (Baytril 2 mg tablets). After recovery, rats were acclimated extensively to the recording set up as well as to handling procedures.

### 2.3. EEG recording

Rats were briefly anesthetized (<2 min) with isoflurane (5% in 95% oxygen) and their head mounts were connected to a preamplifier and recording tethers. Once the righting reflex was fully restored (<5 min), they were placed in Plexiglas recording chambers and connected to the recording setup through a shielded cable carrying a head stage preamplifier (10X) (Pinnacle Technology, Lawrence, KS). EEG signals were routed through an analog adapter to a differential amplifier (100X; DP-304A, Warner Instruments Inc). Thus, the EEG signal was amplified a thousand-fold. Auditory stimuli were ∼ 1 ms square waves generated by Signal v7.0x software (Cambridge Electronics Design (CED), Cambridge, UK) and delivered through the 4-Ohm house speakers (DROK, B00LSEVA8I), fixed at the center of the ceiling of the recording chambers. Auditory stimuli were presented during the course of a 4 s epoch and included either a two-second-long click train at 20 Hz or a one-second-long click train at 40 Hz, both at ∼ 65 dB. In both cases, stimulus train began 1-s after epoch onset. Since 20 Hz and 40 Hz train stimuli were randomized, inter-train interval also varied randomly between 2 and 3 s. EEG was acquired by the data acquisition system (CED Micro 1401-3) as sweeps, time-locked to the 4s epochs. ASSR was averaged from a maximum of 75 trials. Robust harmonics were seen for both 20 Hz and 40 Hz ASSR. However, in this report, we only discuss 20 Hz ASSR and its first three SSHR.

### 2.4. EEG Data analysis

The use of head stage amplification greatly reduced movement artifacts; however, chewing-associated artifacts were noted occasionally. Trials were excluded if an artifact coincided with the auditory stimulus. Thus, typically <10% of the 75 trials were excluded per subject. Preliminary data analysis was performed using Signal v7.01 software. Artifact free trials were averaged for each rat. The averaged data were digitally band-pass filtered (center frequency ±2.5 Hz; 2nd order Butterworth, zero-phase shift, infinite impulse response (IIR) filter). Filtered (as well as unfiltered) averages for each rat showed clear stimulus-associated entrainment. Filtered ASSR or SSHR from individual rats were overlaid to examine how synchrony evolved across the group. Time-frequency analysis (TFA) was performed in the EMSE data editor software v5.6.1 (Cortech solutions, Wilmington, NC). Single trial raw data files from signal software v7.1 were imported as text into EMSE v5.x software (Cortech Solutions Inc., Wilmington, NC). Analysis was performed after Morlet transformation to measure stimulus-induced inter-trial phase coherence (ITPC) and evoked power. The response period 0.2-2 s from train onset was selected for steady state ITC and evoked power comparisons. The same metrics were also computed for a 0.5 s period preceding the train stimulus.

### 2.5. Statistical analysis

Data are expressed as group mean and SEM. A two-way analysis of variance (ANOVA) with repeated measures (mixed-effects model; GraphPad Prism, v9.1) was used. Frequencies and dose levels were designated as row and column factors, respectively. Quantile-quantile (q-q) plots were used to ensure data normality. In case the data were non-normal, they were log transformed first, linearity was confirmed through q-q plotting and only then, parametric statistics were applied. Greenhouse-Geisser correction was implemented without assuming sphericity. Dunnett’s post hoc analyses were used to identify effective doses with vehicle group as comparator.

### 2.6. Pharmacology

MK801 maleate (Sigma-Aldrich) was dissolved in sterile normal saline; free base was used to calculate doses. Injection volume was 1 ml/kg, sc; injections were administered around the same time when the recording tethers were attached to head mounts. The same set of rats were used in a randomized, blocked design with crossover. Post-treatment, spontaneous EEG was recorded for a few minutes to confirm signal quality followed by event related potentials in the presence of auditory clicks at 30, 60, and 90 minutes after dosing. Thus, a pretreatment baseline was not part of this protocol. A washout period of at least 3 days was allowed between dosing.

## 3. Results

### 3.1. Evoked response, spectral peaks and SNR

In vehicle treated rats, grand averaged EEG showed a large N1 response followed by smaller evoked responses preceded by discrete stimulus artifacts, reflecting individual clicks (Figure 1 and inset). Preliminary analysis showed little difference between any of the post-drug time points. Therefore, we focused only on the ∼ 60 min data; moreover, this time corresponds to the C_max_ of MK801 in rats (Wegener et al., 2011). The power spectrum of the evoked response for the duration of the click train showed several discrete peaks, corresponding to the 20 Hz ASSR and its SSHR multiples as marked in Figure 2 (dark line). Note that this pattern contrasted with the power spectrum of the EEG trace for the duration immediately preceding the stimulus train where such peaks were absent (gray line, Figure 2). Both spectra displayed an exponential decay in signal power (Figure 2). This decay was also reflected in ASSR and SSHR peak powers. In the entire power spectrum, there were numerous harmonics of 20 Hz ASSR (not shown). We chose to analyze the first three resolved harmonics which also happened to be the largest. The peak power from these 3 harmonics combined constituted 156% of the peak ASSR power; the later harmonics were substantially smaller (amounting individually to ≤12% of peak ASSR; data not shown). Yet, the signal to noise ratio (SNR) was markedly better at SSHR, relative to ASSR. For example, while the SNR was 3.86 for 20 Hz ASSR, it was 10.28, 21.52 and 45.17 at 40, 60 and 80 Hz SSHR, respectively.

**Figure 1.**
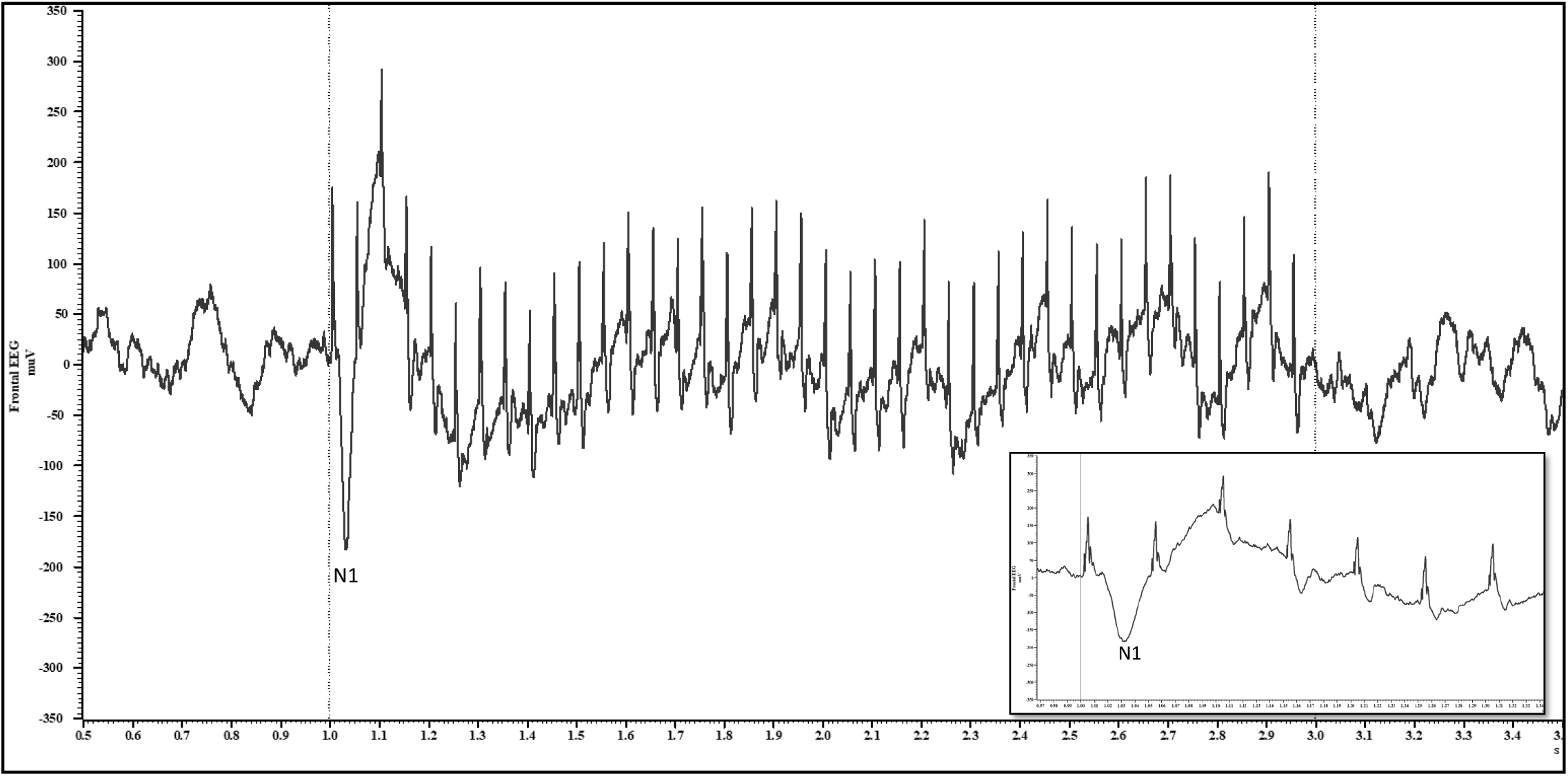
The grand average of the 20 Hz ASSR from vehicle-administered rats (N=12), 60 min after dosing. Note that the spike-like upward deflections are stimulus artifacts while each of these is followed by a slower negative going waveform beginning with the very prominent N1 response. Inset shows the first few evoked responses, highlighting a large N1 response followed by smaller and apparently gated negative waves. Train onset and off set are marked by dashed vertical lines.

**Figure 2.**
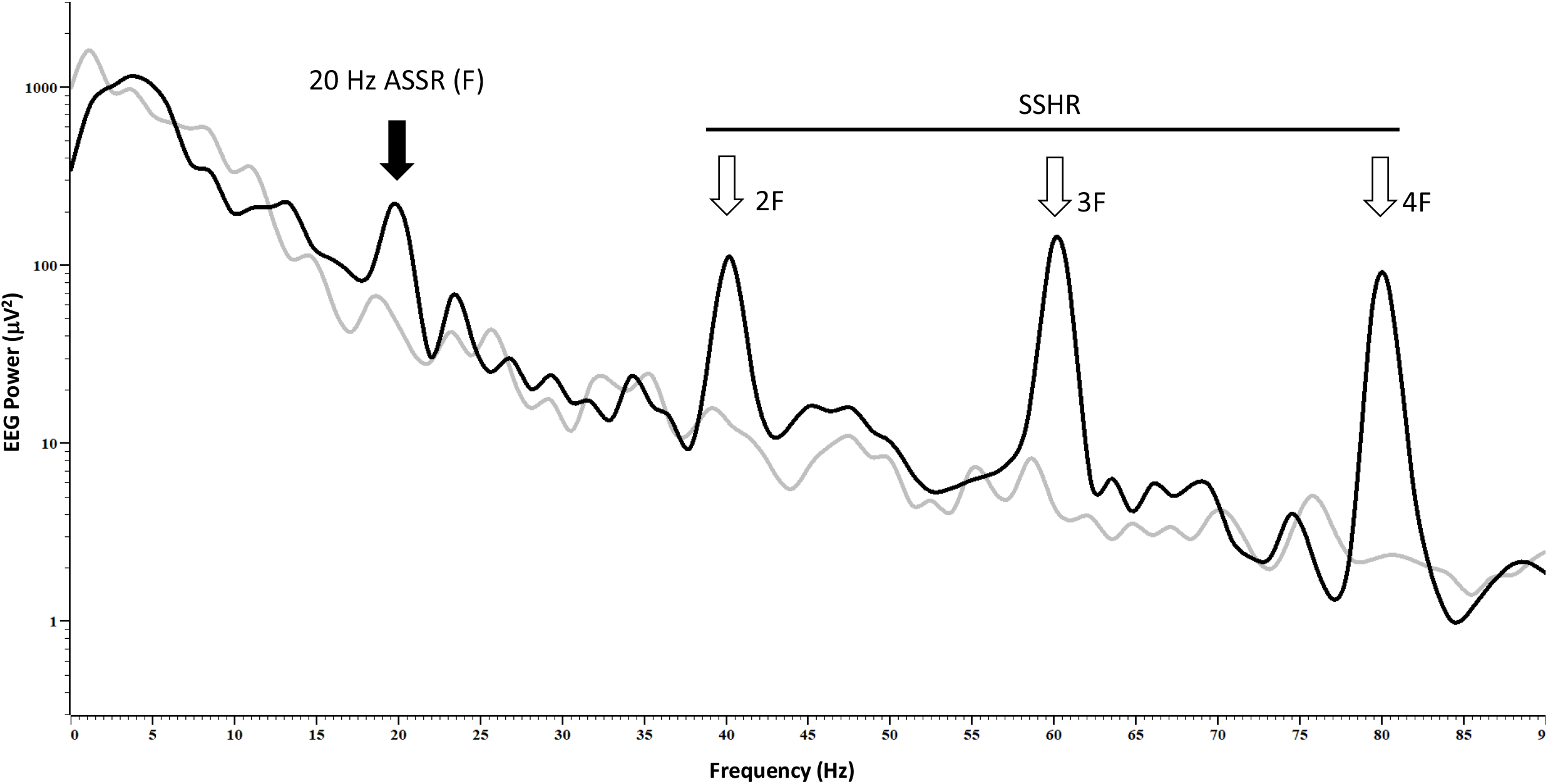
Grand averaged power spectrum of the 20 Hz ASSR from vehicle-dosed rats 60 min post-administration. The power spectrum corresponds to a 0.9 s long ASSR segment after the steady state response was established (0.3-1.2 s post stim; dark line). Grey line corresponds to a period of 0.9 s before stim onset. A Hanning window with an FFT size of 8192 was used for both spectra.

### 3.2. ITPC in vehicle treated rats

While the evoked power at 20 Hz ASSR was marginally higher than the evoked power at the first three harmonics, ITPC showed a contrasting trend. ASSR at 20 Hz had relatively weak ITPC and it progressively increased from 40-80 Hz. These data are summarized in Figure 3.

**Figure 3.**
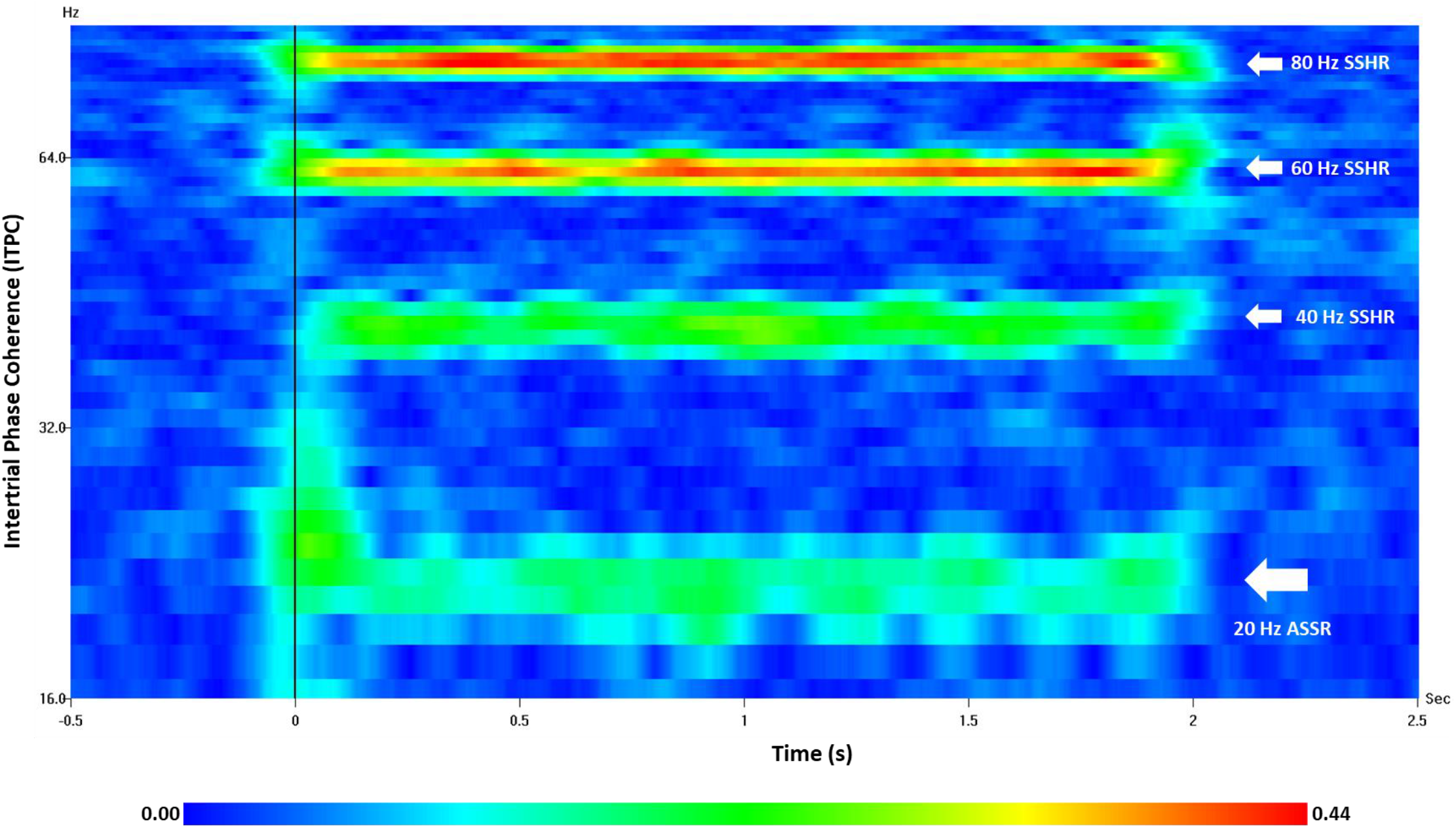
Intertrial phase coherence of 20 Hz ASSR from vehicle-dosed rats (N=12) and the three successive SSHR. Note the higher phase-locking at 60 and 80 Hz harmonics.

### 3.3. Temporal evolution of neural synchrony

To examine how ASSR and SSHR evolved over time, narrow band filtered (center frequency ±2.5 Hz) EEG traces were overlaid from a group of 12 rats at F, 2F, 3F and 4F (Figure 4). Visual evaluation indicated that synchrony evolved variably across ASSR and SSHR; whereas synchrony at 60 Hz became apparent by ∼ 20 ms from train onset, 20 and 80 Hz oscillations took ∼ 60 and ∼ 80 ms for entrainment, respectively. Entrainment at 40 Hz SSHR took the longest time, about 180 ms from train onset, before a clear group-wide synchrony became apparent (Figure 3). Indeed, relative to the ITPC measured over 100 ms pre-stimulus vs. 100 ms post-stimulus, no significance was noted at 40 Hz SSHR (p>0.05, pared t-test). As expected, there was a significant increase in ITPC post-stimulus at all other frequencies (p>0.01; data not shown).

**Figure 4.**
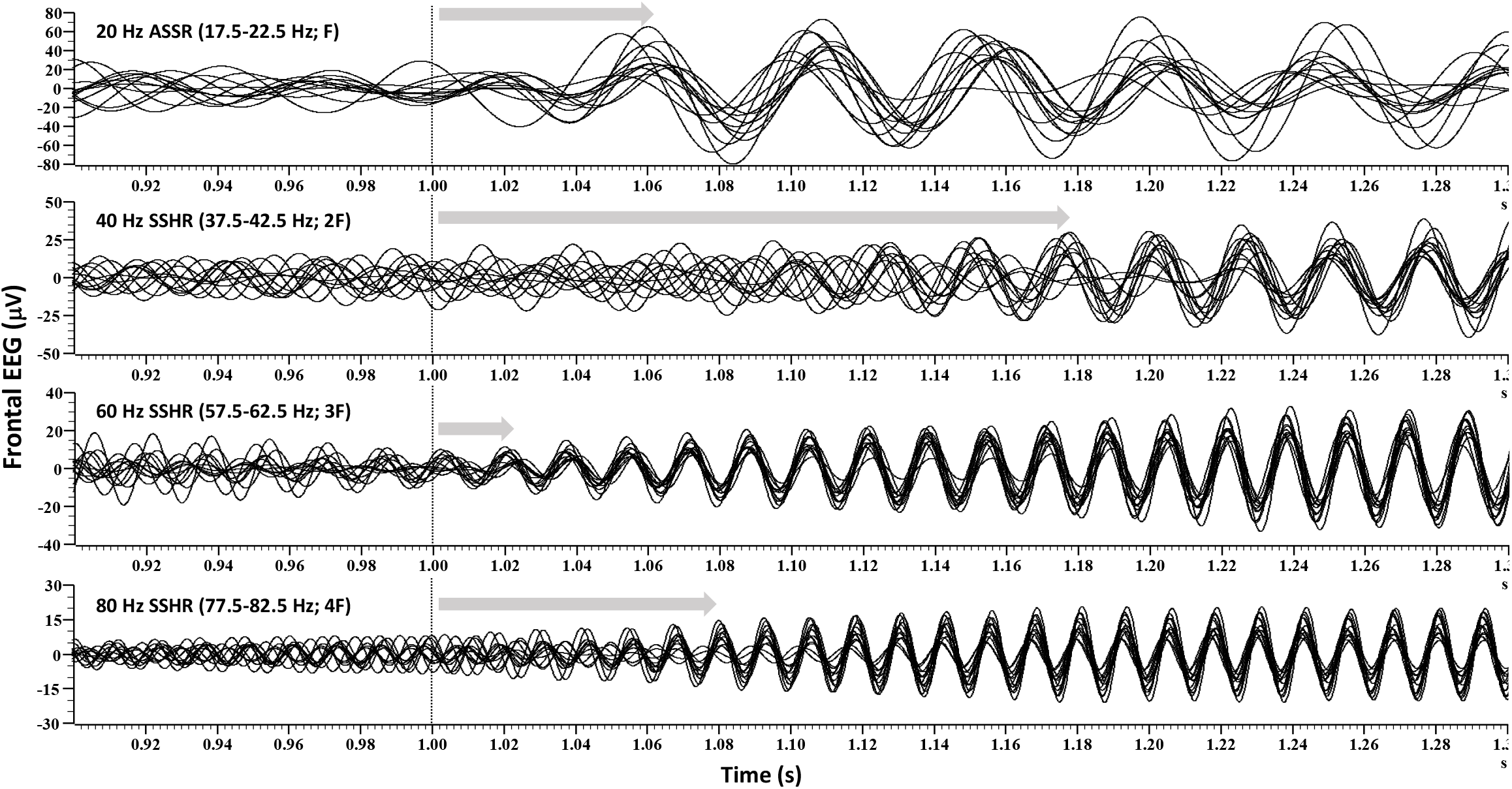
Narrow band-pass filtered ASSR and SSHR traces from vehicle treated rats (N=12) are overlaid to indicate how entrainment evolved at each frequency. Length of arrow indicates the time after stimulus onset, when entrainment became apparent across the group. Vertical line marks stimulus onset.

### 3.4. Differential effects of MK801 on ASSR and SSHR

#### ITPC

To characterize the ASSR and SSHR, we examined the intertrial phase coherence (ITPC) in a narrow frequency band centered around 20 Hz (±2.5 Hz), as well as across the first three harmonics corresponding to 40 Hz, 60 Hz and 80 Hz (±2.5 Hz at each frequency). These values were assessed after vehicle injection and after administration of three active dose levels of the NMDA antagonist, MK801. The results are summarized in Figure 5, left panel. Two-way ANOVA with repeated measures showed a strong effect of frequency [p<0.0001; F (2.351, 25.86) = 105.3; epsilon = 0.7836], dose [p<0.0001; F (1.522, 16.74) = 59.13; epsilon = 0.5074] and a frequency by dose interaction [p<0.0001; F (3.872, 42.59) = 17.75; epsilon = 0.4302]. The mean ITPC at 20 Hz was 0.17 after vehicle injection and was significantly decreased after the 0.025 mg/kg dose only (p<0.05). Higher MK801 doses had no significant effect on ITPC (p>0.05). In contrast, strong and dose-dependent reductions in ITPC were noted across all three harmonics, with p values ranging from ≤0.001 to 0.0001 (Figure 5, left panel). Moreover, in the vehicle condition, from visual inspection of the data, it was apparent that the group synchrony at SSHR was stronger than at the ASSR (Figure 3).

**Figure 5.**
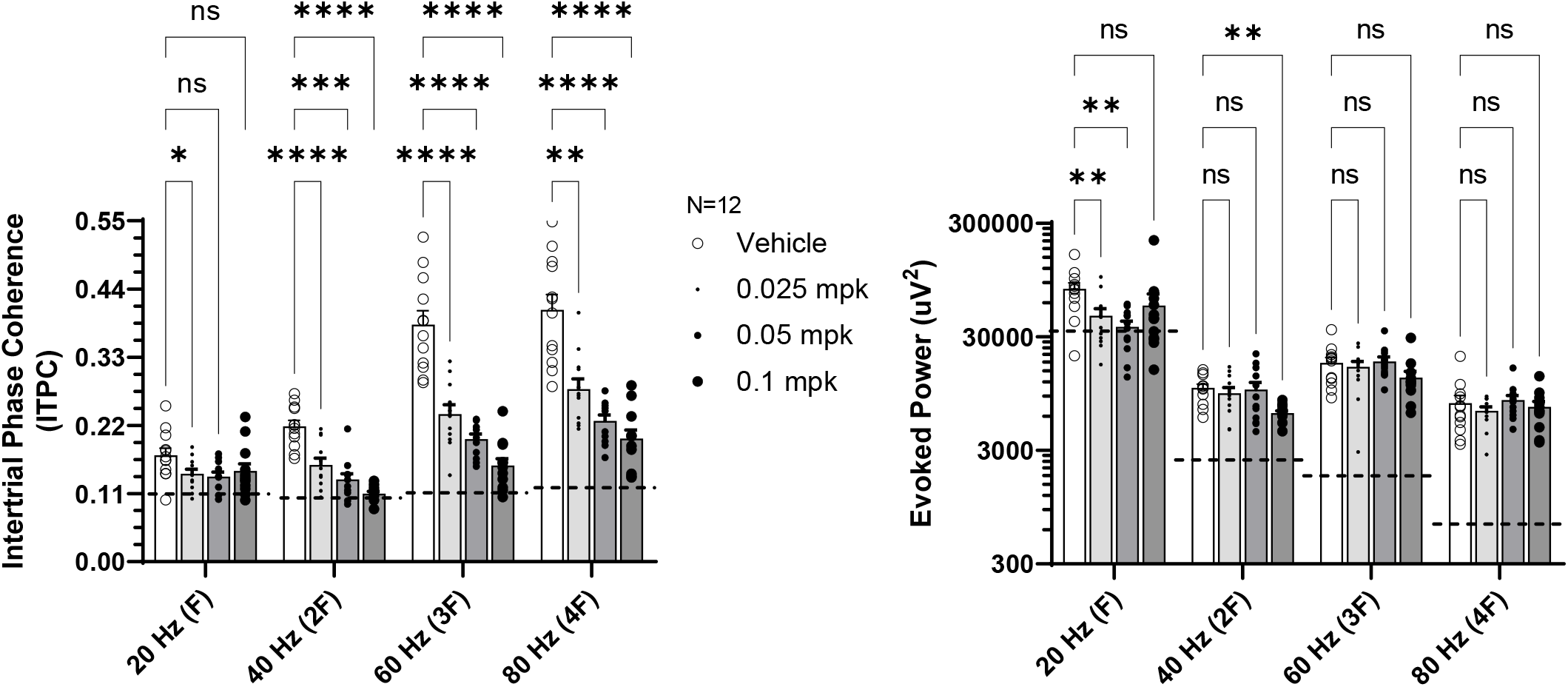
Compiled data representing the mean intertrial phase coherence (left panel) ± SEM and evoked power (right panel) ± SEM of ASSR and SSHR after dosing with vehicle or MK801. The horizontal dashed lines indicate the “floor” for this assay. Note strong dose-dependent reductions only for the SSHR. In contrast, evoked power was reduced by MK801 at the 20 Hz ASSR and 40 Hz SSHR only. ^*^/^**^/^***^ indicate p<0.05/0.01/0.001, respectively.

We next evaluated how well the variance in ITPC at ASSR predicted the variance in harmonics. The R^2^ values from this analysis indicated that while statistically significant, they were quite modest, suggesting that only a small portion of variance in SSHR could be predicted by the variance in ASSR. The R^2^ values and corresponding p values for ITPC were as follows: 20 Hz ASSR vs 40 Hz SSHR (0.16; p=0.0044); 20 Hz ASSR vs 60 Hz SSHR (0.23; p=0.0005); 20 Hz ASSR vs 80 Hz SSHR (0.23; p=0.0006).

### 3.5. Evoked power

We next examined the evoked power during click train presentation. Two-way ANOVA revealed a strong frequency effect [p<0.0001; F (2.871, 31.58) = 204.7; epsilon = 0.9570], and significant frequency by dose interaction [p=0.0045; F (4.042, 44.47) 4.363; epsilon = 0.4492], but no main dose effect (p>0.05). Posthoc comparisons between drug level and vehicle at each frequency showed that 20 Hz ASSR evoked power was attenuated by MK801 at 0.025 and 0.05 mg/kg but not at 0.1 mg/kg. In contrast, a strong attenuation of 40 Hz SSHR evoked power was noted at the 0.1 mg/kg dose only, relative to vehicle. Lastly, SSHR evoked power at 60 and 80 Hz were unaffected by MK801. These results are summarized in Figure 5, right panel.

### 3.6. Prestimulus ITPC and signal power

To determine the minimum value of ITPC in our experiments and to evaluate the effect of drug administration on spontaneous narrow band signal power, we calculated ITPC and narrow band signal power during a period of 0.5 s preceding click train onset in single trials. The ITPC ranged between 0.10-0.12 across frequencies, suggesting that this is the working limit for this measure in our assay. These levels are indicated in figure 4 as dashed lines across frequencies and treatments. Moreover, there were no frequency, dose or frequency by dose effects (p<0.05) (data not shown). In contrast to ITPC, there were strong effects on the total signal power for frequency [p<0.0001; F (1.946, 21.40) = 2609; epsilon = 0.6486], dose [p<0.0001; F (1.842, 20.26) = 43.85; epsilon = 0.6141] and a frequency by dose interaction [p<0.0001; F (3.019, 33.20) = 65.60; epsilon = 0.3354] (Figure 6). Moreover, the direction of change due to dose was informative across frequencies; while a strong suppressive effect of MK801 was noted on the prestimulus beta band power (p-values ranged from <0.01-0.0001), dose-dependent increases were noted at the three harmonics tested (p values ranged from <0.001-0.0001) (data not shown). Lastly no significant coefficients of determination (R^2^) were noted between prestimulus ITPC of 20 Hz ASSR and SSHR at 40, 60 or 80 Hz (data not shown).

**Figure 6.**
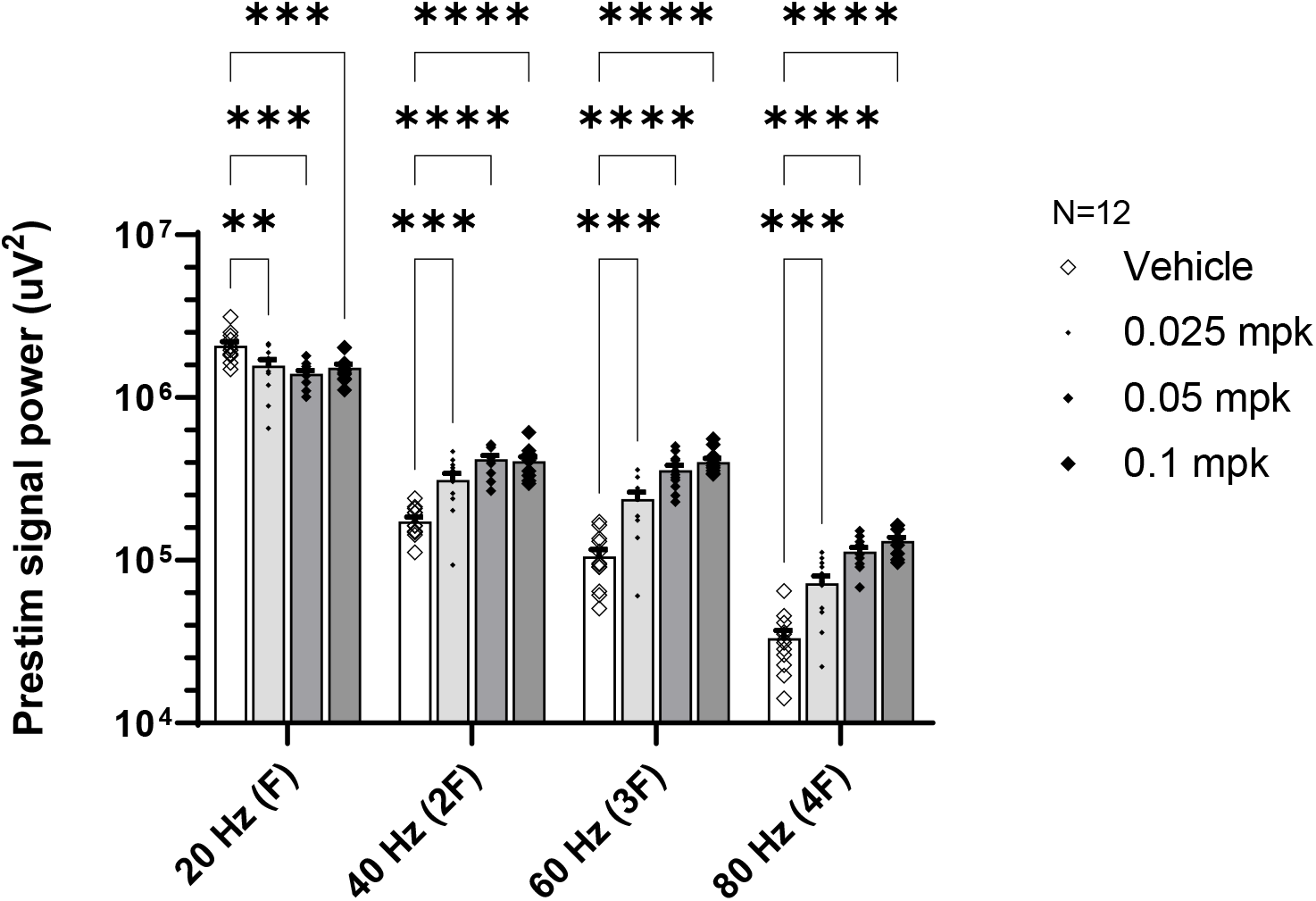
Compiled narrow band (center frequency±2.5 Hz) signal power during the prestimulus condition. MK801 dose-dependently reduced beta band power while strongly and dose-dependently increasing background gamma activity. ^**^/^***^/^****^ indicate p<0.01/0.001/0.0001, respectively.

## 4. Discussion

Periodic and complex stimuli like a train of clicks carry signal power not only at the driving frequency F but also at its integer multiples (n^*^F), called harmonics (Bizley and Walker, 2010; Coffey et al., 2019). Since the sound vibrations are at distinct frequencies, they activate different regions of the cochlear basilar membrane and eventually route to tonotopically represented regions in the cortex. The literature on click train mediated response has generally maintained its focus on the driving frequency response and not on harmonics (Puvvada et al., 2018; Tsuchimoto et al., 2011). However, a few reports have indicated that in schizophrenia patients, 20 Hz ASSR was accompanied by a significant and selective reduction in the second harmonic at 40 Hz (Kwon et al., 1999; Spencer et al., 2008). Given the important role of harmonics in auditory processing across species (Bizley and Walker, 2010; Bondar and Shubina, 2018; Obleser et al., 2008; Plack et al., 2006; Popham et al., 2018), and the unique advantage associated with steady state response as a high SNR neural synchrony measure (Norcia et al., 2015), we were interested in exploring the idea that SSHR could be a non-redundant biomarker for pharmacological intervention. Once thoroughly validated, such measures are critical for CNS drug development, an area that currently lacks reliable disease-relevant functional biomarkers critical for confirming target engagement, enable dose selection and accomplish patient stratification (Light and Swerdlow, 2020; Luck et al., 2011; O’Donnell et al., 2019).

Many neurophysiological deficits seen in schizophrenia patients are acutely mimicked in animal species by the administration of NMDA antagonists (Bristow et al., 2016; Curic et al., 2019; Ehrlichman et al., 2009; Gunduz-Bruce, 2009; Leishman et al., 2015; Sivarao et al., 2014). In this study, we tested if the high affinity, open NMDA channel blocker, MK801, would disrupt the 40 Hz harmonic of the 20 Hz ASSR, modeling what has been reported previously in schizophrenia patients (Kwon et al., 1999; Spencer et al., 2008; Vierling-Claassen et al., 2008). Moreover, given the contrasting modulation of 20 Hz ASSR vs its SSHR at 40 Hz, the non-redundant role of harmonics in sound processing (Bizley and Walker, 2010; Driesen et al., 2013; Oxenham, 2012) and their separable localization relative to the steady state response (Hamm et al., 2012; Heinrichs-Graham and Wilson, 2012), we hypothesized that ASSR and SSHR would have distinct characteristics, e.g. neural synchrony metrics (Bondar and Shubina, 2018) and/or pharmacological responsivity. In the present analysis, we characterized the first three harmonics, which were well resolved as well as the largest among all harmonics, in terms of their peak power (Figure 2). It however needs to be acknowledged that the findings discussed below were from female rats only. In future studies, these observations need to be replicated in adult male rats for generalizability.

In support of our hypothesis, analyses revealed several differences between ASSR and SSHR, as discussed below. For example, the power of SSHR tended to be lower than the power of ASSR at 20 Hz. Since the train-mediated oscillations ride on top of on-going EEG whose power spectrum exhibits an inverse relationship with frequency, it was not surprising that the higher frequency SSHR had lower power. However, perhaps for the same reason, the higher harmonics had a stronger signal-to-noise ratio, compared to the ASSR. In human literature, however, such data appear variable; for example, relative to 40 Hz SSHR, evoked power at 20 Hz ASSR was found to be lower (Hamm et al., 2011; Ross et al., 2000), similar (Kwon et al., 1999; Spencer et al., 2008) or larger (Tsuchimoto et al., 2011). Perhaps reflecting this diversity, relative to gamma frequency ASSR, ASSR at 20 Hz has been shown to be more variable in a test-retest evaluation (Roach et al., 2019).

Unlike evoked power, ITPC was the weakest at the 20 Hz ASSR and progressively increased with the harmonic number. As a result, the phase-locking exhibited by the SSHR was markedly higher than the phase-locking at the ASSR. This difference may be because the SSHRs were in the gamma frequency range and may reflect the resonance properties of frontal networks that preferentially entrain to gamma frequencies over lower frequencies (Ferrarelli et al., 2012; Tremblay et al., 2019). Notably, scalp recorded human ASSR studies show that the signal power is highest ∼ 40 Hz relative to other frequencies, either in the beta range or in the high-gamma range (Galambos et al., 1981; Picton et al., 2003). In fact, this is what led early investigators to surmise that ∼ 40 Hz represents a resonant frequency of neocortical networks. It is likely that in rodent neocortex too, stimuli in the gamma range of 30-100 Hz elicit the most resonance. However, the precise frequency for peak resonance may differ based on the exact region of recording, species and perhaps even the technique used. For example, recordings from rat temporal cortex have shown that 40 Hz stimuli elicit the most signal power (Conti et al., 1999; Kozono et al., 2019). On the other hand, Kuwada et al. demonstrated a peak ASSR response at 62 Hz in rabbits (Kuwada et al., 2002). Recently, Nakamura et al. examined a range of frequencies in macaque monkeys and found that click stimuli at ∼ 80 Hz and not ∼ 40 Hz, elicited the maximum signal power and phase locking (Nakamura et al., 2022).

We examined whether the phase locking pattern in harmonics is a simple reflection of the phase-locking at the ASSR. Looking at the frequency-wise overlay of the traces, as shown in Figure 4, it is apparent that it took about 60 ms from stimulus onset for nearly all rats to show a 20 Hz (driving frequency) oscillatory pattern, while it only took 20 ms for the 60 Hz harmonic to entrain. This at first seemed logical since the 60 Hz harmonic wave has a period that is one-third of the 20 Hz period. However, the 80 Hz harmonic with a 12.5 ms period took almost 80 ms to emerge as a coherent oscillation (Figure 4). Lastly, the 40 Hz harmonic took the longest time to emerge, with group-wide synchrony establishing ∼ 180 ms from stim onset. This delayed onset of 40 Hz SSHR parallels the slow emergence of the 40 Hz ASSR in humans (Ross et al., 2002), as well as in rodent frontal cortex, as reported recently (Ummear Raza et al., 2023). Thus, it is tempting to speculate that the 40 Hz harmonic may engage the same network or a subset of the network activated by a 40 Hz driving stimulus.

Given the differences discussed above, we assessed the correlations of synchrony measures at 20 Hz with those at the three harmonics. As discussed above, there were statistically significant but modest-sized correlations between 20 Hz ASSR and 40 Hz SSHR for ITPC and evoked power. However, no significant correlations were seen between ASSR and SSHR at 60 Hz or 80 Hz for ITPC or evoked power (data not shown).

We next examined how treatment with the high affinity NMDA open channel blocker affected the ASSR and harmonics. Effects of MK801 on ITPC at 20 Hz ASSR were inconsistent: whereas the lowest dose (0.025 mg/kg) was associated with a small but significant reduction in ITPC, higher doses did not affect this measure. In contrast, a robust and dose-dependent disruption of ITPC was noted at all three harmonics (Figure 4, left panel). Evoked power at 20 Hz ASSR was also inconsistently affected by MK801: it was reduced at the lower two doses but was unaffected at the highest dose of 0.1 mg/kg (Figure 4, right panel). In contrast, only the highest MK801 dose suppressed the SSHR at 40 Hz relative to vehicle. Evoked power at higher harmonics of 60 and 80 Hz were unaffected by MK801 (Figure 4, right panel). Thus, pharmacologic modulation of harmonics by MK801 was frequency specific and differentiated relative to its effects on the ASSR.

In the prestimulus condition, we computed ITPC as well as the total power. Since there is no stimulation during this time frame, the ITPC simply indicates what the “floor” is for this measurement. Although the ITPC range was smaller for the ASSR, it still allowed ample room for pharmacological modulation. Computed total power during the same period showed that baseline narrow band beta and gamma were differentially modulated by MK801: beta power (∼ 20 Hz) was reduced while gamma power was robustly increased, in line with the known effects of NMDA antagonists on background gamma and E/I balance (Pinault, 2008; Saunders et al., 2012).

In summary, click trains elicit entrainment not only at the driving frequency but also at higher harmonics. In the context of the literature on ASSR in neuropsychiatry, these responses have not been the focus of many studies. Given the importance of harmonics in acoustic processing and their non-redundancy relative to the response at F, and given the signal to noise advantage of narrow band entrainment in general, we hypothesized that it would be possible to record harmonics, and that they would exhibit distinct characteristics including pharmacologic sensitivity, relative to the response at the driving frequency. Indeed, using 20 Hz ASSR and its three successive harmonics, we demonstrate strong and distinct modulation of SSHR, relative to the ASSR. Such distinct characteristics underscore the fact that the SSHR is a novel pharmacodynamic neural circuit biomarker that could provide additional value to conventional ASSR studies. As discussed above, even when the ASSR is non-responsive to a pharmacological intervention, the SSHR may offer a new opportunity for demonstrating functional engagement with excellent sensitivity and SNR. Thus, SSHR appears to be a novel biomarker that can be exploited for future drug development.

## Abbreviations

ASSR: auditory steady state response
SSHR: steady state harmonic response

